# RAFTER: a releasing factor tethered RNA editing system for quantifying mRNA translation

**DOI:** 10.64898/2025.12.12.693956

**Authors:** Zhiyuan Sun, Peng Dong, Huanyu Yan, Xiaozhen Wen, Yanping Li, Feifei Jiang, Liang Fang, Xi Wang, Wei Chen

## Abstract

Measurement of mRNA translational efficiency has heavily relied on quantifying the amount of polysome-associated RNAs which would not always reflect the efficient translation. To address this, we developed RAFTER, which combined RNA editing enzyme with translation releasing factor to selectively label successfully translated RNAs. The translational efficiency estimated based on RAFTER correlated well with that by Ribo-seq and Polysome-seq, but with greatly simplified workflow, which enables it for single-cell analysis.

## Main

Translational regulation plays a fundamental role in shaping cellular physiology by modulating protein synthesis^1^. Dysregulation of mRNA translation has been implicated in numerous diseases, including cancer, neurodegenerative disorders, and developmental abnormalities^2-4^. Therefore, precise and reliable quantification of translational efficiency are crucial for understanding the intricate mechanisms underlying gene expression control. Over the years, several techniques have been developed to measure translational efficiency, each with its own strengths and limitations. Polysome profiling, a classical technique, enables the separation of mRNA molecules bound to varying numbers of ribosomes, with higher numbers indicating more active translation^5^. Ribosome footprinting, an alternative tool, involves deep sequencing of ribosome-protected mRNA fragments, which allows determination of the precise positions of ribosomes on individual transcripts^6^. Although these two techniques have greatly advanced our understanding of translational regulation, high ribosome association with mRNAs does not necessarily indicate high translation efficiency, which also depends on the translation elongation rate^7^. Moreover, both methods generally require large amounts of input material, which limits their application to rare or limited samples. To address these challenges, RNA editing-based strategies have recently emerged as powerful tools for measuring ribosome-RNA interactions^8, 9^. By coupling deaminase enzymes with small ribosome subunit proteins, these approaches enable in situ labeling of ribosome-associated RNAs and can even be applied to identify ribosome-RNA interactions at single-cell resolution^9-11^. However, the association between ribosomes and mRNAs does not necessarily indicate successful translation. To overcome this limitation, we leveraged the translation-releasing factor, which integrates into the ribosome complex specifically during the translation termination stage.

To develop a translation releasing factor-based RNA editing method for quantifying mRNA translation rates, we first analyzed the protein components of the termination complex. The mammalian termination complex is composed of two release factors: eRF1 and eRF3. eRF1 recognizes stop codons and stabilizes the binding of GTP to eRF3^12^, whereas eRF3 is a GTPase that hydrolyzes GTP to stimulate peptide release by eRF1^13^. The cryo-electron microscopy structure of the mammalian termination complex revealed that the N-terminal region of eRF1 reaches deep into the decoding center, while eRF3 is located on the surface of the complex^14^. Therefore, to mitigate potential structural interference from the release factor-deaminase fusion protein, we selected eRF3 as the candidate. In mammals, there are two distinct eRF3 genes: the widely expressed eRF3a (GSPT1) and the tissue-specific eRF3b (GSPT2)^14^. To enable broad applicability, we chose eRF3a and employed an eRF3a-ADAR2dd fusion protein, which was expressed in a doxycycline-induced manner (Fig. 1a). The method is named RAFTER (ReleAsing Factor Tethered Editing of RNA).

**Figure 1.**
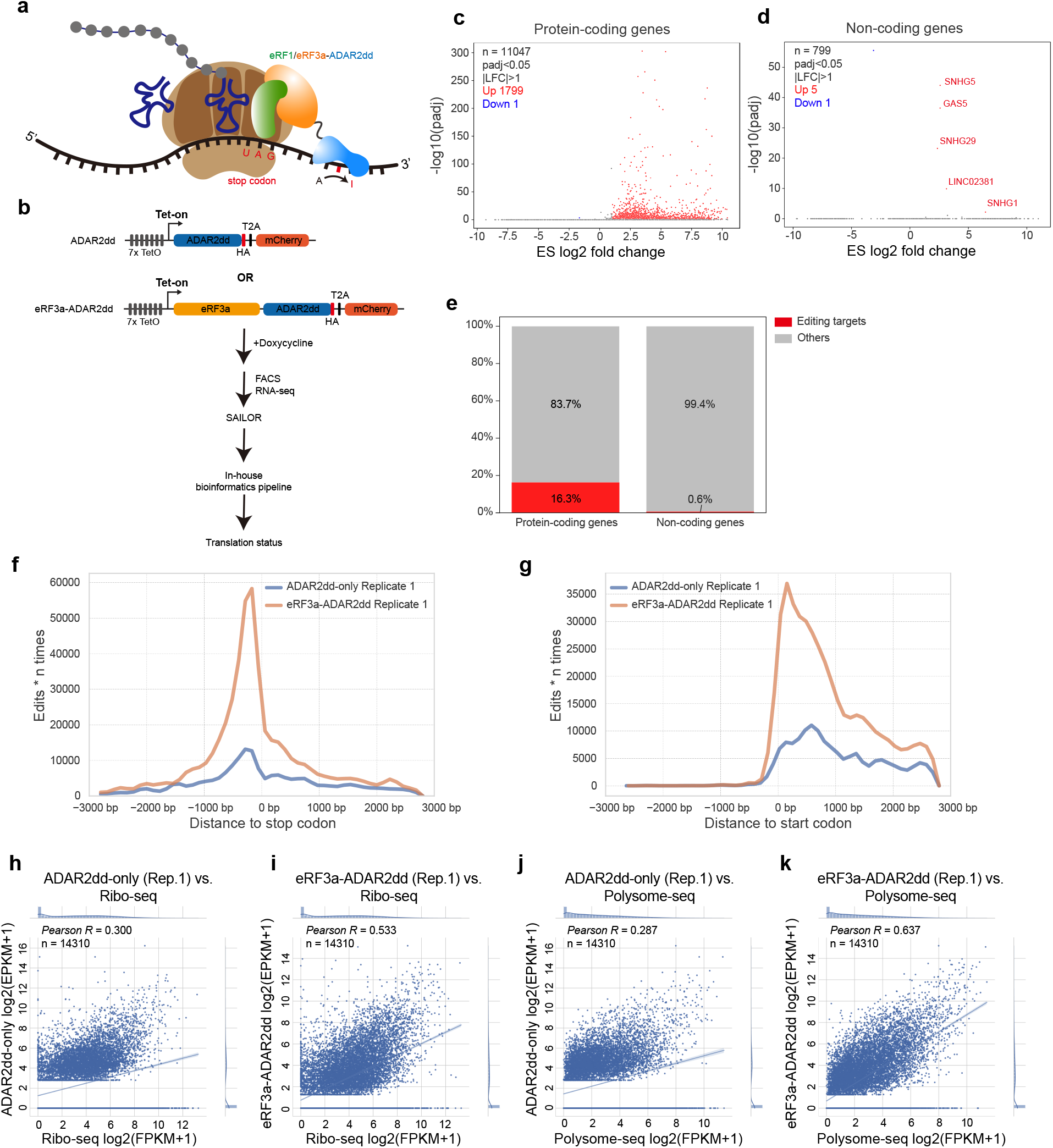
Establishment of RAFTER. **a**, Schematic representation of the RAFTER method. **b**, Experimental and analytical workflow of RAFTER. **c**, Volcano plots comparing the editing score (ES) of protein-coding genes between cells expressing eRF3a-ADAR2dd and ADAR2dd-only. The X and Y axes represent log2 fold change (LFC) and −log10(p-value), respectively. Statistical significance was calculated using two-sided Fisher’s exact test followed by Benjamini-Hochberg correction. **d**, Volcano plots comparing the editing score (ES) of non-coding genes between cells expressing eRF3a-ADAR2dd and ADAR2dd-only. The X and Y axes represent log2 fold change (LFC) and −log10(p-value), respectively. Statistical significance was calculated using two-sided Fisher’s exact test followed by Benjamini-Hochberg correction. **e**, Bar chart showing the percentages of editing targets identified in protein-coding and non-coding genes, respectively. **f-g**, Metaplots showing the distribution of confident edit sites (≥ 0.65 confidence score) in the flanking regions of stop codons (f) and start codons (g). **h-k**, Scatter plots showing the correlation between RNA editing levels and RNA translation levels. **h**, ADAR2dd-only (replicate 1) vs. Ribo-seq. **i**, eRF3a-ADAR2dd (replicate 1) vs. Ribo-seq. **j**, ADAR2dd-only (replicate 1) vs. Polysome-seq. **k**, eRF3a-ADAR2dd (replicate 1) vs. Polysome-seq. The number of genes (n) and *Pearson* correlation coefficient (*R*) are indicated in the top left corner of each panel.

In RAFTER, the cells expressing ADAR2dd-only proteins served as background editing controls. After 48 hours of doxycycline induction, cells exhibiting equilibrated exogenous gene expression, as determined by mCherry intensity using fluorescence-activated cell sorting (FACS), were collected for RNA-seq (Fig. 1b and Supplementary Figure 1a-c). After identifying confident edit sites using SAILOR^15^, we calculated the editing score (ES) for each gene based on edit sites with a confidence value ≥0.65 in each sample, as previously described^16^. Using this strategy, 1799 annotated protein-coding genes were identified as editing targets, which have significantly higher ES in eRF3a-ADAR2dd expressing cells than in ADAR2dd-only controls (Fig. 1c). In contrast, only five annotated long non-coding RNAs (lncRNAs) were identified as editing targets (Fig. 1d). As expected, the proportion of identified targets were significantly higher among annotated protein-coding genes than among lncRNAs (Fig. 1e). Interestingly, four of the five non-coding genes with higher ES in eRF3a-ADAR2dd expressing cells (SNHG1, SNHG5, GAS5, and SNHG29) have previously been reported as translated, supported by Ribo-seq and/or mass spectrometry^17-19^. These together indicate the specificity and sensitivity of our method for detecting translating RNAs.

Since the termination complex binds to mRNAs when the ribosome encounters a stop codon, we next evaluated whether RAFTER-mediated RNA editing could accurately reflect the association between the termination complex and mRNA by plotting the distribution of confident edited reads (see Methods). As expected, eRF3a-ADAR2dd editing resulted in prominent peaks around the stop codon of protein-coding genes, while the control protein only generated much sparser edit events (Fig 1f, Supplementary Figure 2a). Meanwhile, we noticed that eRF3a-ADAR2dd also generated editing peaks around the start codon, with limited extension into the 5’ UTR of mRNAs (Fig. 1g, Supplementary Figure 2b). This indicates a potential interaction between the start and stop codons of mRNAs, which may represent an additional loop structure, distinct from the known 5’ cap-3’ poly(A)-tail interaction^20^, during mRNA translation.

**Figure 2.**
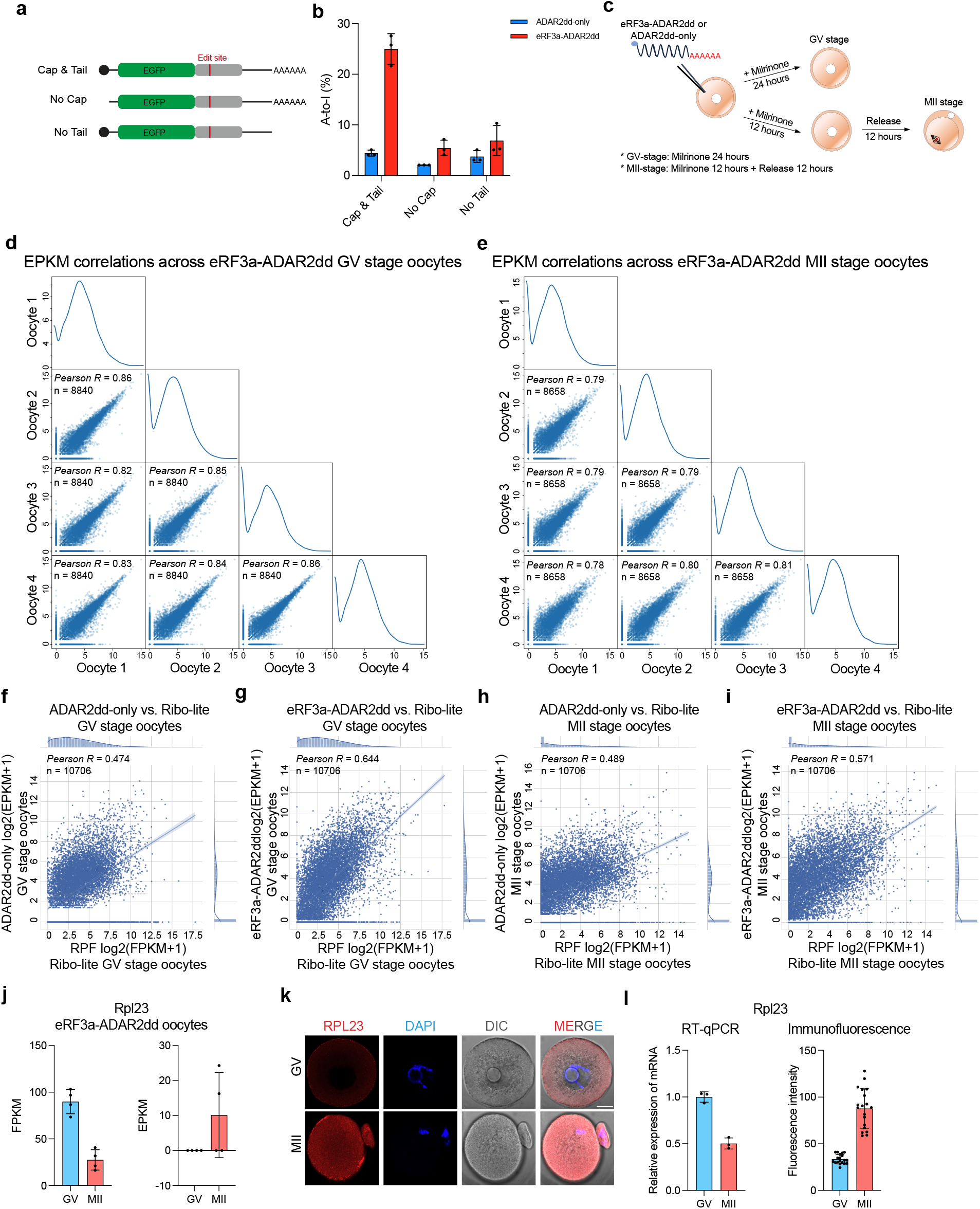
RAFTER captures bona fide translation and translation landscape in single mouse oocyte. **a**, Design of different reporter mRNAs. **b**, Bar plots showing the RNA editing level of different reporter mRNAs. n = 3. p (cap & tail) < 0.0001, p (no cap) < 0.1491, p (no tail) < 0.1890. Statistical significance was calculated using ordinary two-way ANOVA (default in Prism 9). **c**, Experimental workflow of applying RAFTER on mouse oocytes. **d**, Scatter plots showing the correlation of EPKM across individual eRF3a-ADAR2dd mRNA-injected GV stage oocytes. The X and Y axes represent log2(EPKM+1). The number of genes (n) and *Pearson* correlation coefficient (*R*) are indicated in the top left corner of each plot. **e**, Scatter plots showing the correlation of EPKM across individual eRF3a-ADAR2dd mRNA-injected MII stage oocytes. The X and Y axes represent log2(EPKM+1). The number of genes (n) and *Pearson* correlation coefficient (*R*) are indicated in the top left corner. **f-i**, Scatter plots showing the correlation between RNA editing levels measured by RAFTER and RNA translation levels measured by the Ribo-lite approach in single mouse oocyte. **f**, in GV stage oocytes, ADAR2dd-only vs. Ribo-lite. **g**, in GV stage oocytes, eRF3a-ADAR2dd vs. Ribo-lite. **h**, in MII stage oocytes, ADAR2dd-only vs. Ribo-lite. **i**, in MII stage oocytes, eRF3a-ADAR2dd vs. Ribo-lite. The X and Y axes represent RPF log2(FPKM+1) and log2(EPKM+1), respectively. The number of genes (n) and *Pearson* correlation coefficient (*R*) are indicated in the top left corner of each panel. **j**, RNA level and editing level of Rpl23 detected in eRF3a-ADAR2dd mRNA-injected oocytes. N = 4. **k-l**, Independent quantification of RNA and protein level of Rpl23 in mouse oocytes. **k**, representative immunofluorescence results showing protein level of Rpl23 in mouse oocyte. Scale bar, 20 μm. **l**, RT-qPCR quantification of RNA level (relative to endogenous 18S rRNA, n = 3) and fluorescence intensity quantification of protein level (from immunofluorescence results, n = 19). p (RT-qPCR) = 0.0031. p (fluorescence intensity) < 0.0001. Statistical significance was calculated using two-sided unpaired t test (default in Prism 9). The experiment was repeated independently for 2 times with similar results.

Since the majority of editing events generated by eRF3a-ADAR2dd were located near the stop and start codons, we defined two windows on each protein-coding gene (−100 bp to +2000 bp relative to the start codon and −1000 bp to +200 bp relative to the stop codon) to more accurately estimate editing efficiency per gene. We normalized the number of edited reads in these regions to the total number of edited reads, yielding edits per kilobase per million total edits (EPKM). As shown in Supplementary Figure 2c and 2d, although EPKM values correlated well between replicates in both induced ADAR2dd-only and eRF3a-ADAR2dd cells, the Pearson correlation coefficient (R) was higher in eRF3a-ADAR2dd expressing cells than in ADAR2dd-only expressing cells, suggesting that RNA editing events generated by ADAR2dd alone are more stochastic than those generated by eRF3a-ADAR2dd. Subsequently, we compared EPKM values obtained using RAFTER with FPKM values obtained using ribosome footprinting (Ribo-seq)^21^ and polysome profiling (Polysome-seq)^22^, two commonly used measures of translational efficiency. In ADAR2dd-only expressing cells, EPKM values were poorly correlated (*Pearson R* around 0.3) with FPKM values from either Ribo-seq (Fig. 1g and Supplementary Figure 2e) or Polysome-seq (Fig. 1i and Supplementary Figure 2g). In contrast, in eRF3a-ADAR2dd expressing cells, these correlations were substantially higher (*Pearson R* around 0.6; Fig. 1i and 1k, Supplementary Figure 2f and 2h), which is comparable to the correlation between Ribo-seq and Polysome-seq (Fig. 1k and Supplementary Figure 2i). Taken together, these results demonstrate that translational efficiency inferred by RAFTER correlates well with measurements from both Ribo-seq and Polysome-seq.

To further investigate whether RAFTER-generated RNA editing reflects bona fide mRNA translation, we designed a reporter system (Fig. 2a). Briefly, an endogenous A site that was robustly edited in eRF3a-ADAR2dd-expressing cells was cloned into the 3’ UTR downstream of the EGFP coding sequence. EGFP–3′ UTR mRNAs were then synthesized by *in vitro* transcription with or without a 5′ cap and with or without a 3′ poly(A) tail. The resulting mRNAs were transfected into doxycycline-induced ADAR2dd-only or eRF3a-ADAR2dd expressing cells for 12 hours. Consistent with previous observation^23, 24^, flow cytometry analysis showed that only capped and polyadenylated mRNA was efficiently translated (Supplementary Figure 3a and 3b). RT-qPCR analysis further indicated that uncapped mRNAs were less stable than capped mRNAs (Supplementary Figure 3b and 3c). Through Sanger sequencing, we observed robust A-to-I editing of the capped and polyadenylated mRNA in eRF3a-ADAR2dd expressing cells, but not in ADAR2dd-only expressing cells (Fig. 2b). Moreover, A-to-I editing levels were substantially lower for mRNAs lacking either the 5′ cap or 3′ poly(A) tail than for the fully modified mRNA in eRF3a-ADAR2dd expressing cells (Fig. 2b). These results together demonstrate that RAFTER reliably reports authentic mRNA translation.

While single-cell ribosome profiling and mass spectrometry have been developed^25-28^, the technical complexity and high instrumentation demand limit their widespread use. In principle, RAFTER offers a feasible alternative for single-cell detection of mRNA translation. To explore this possibility, we applied RAFTER to measure the translational landscape of mouse oocytes during the germinal vesicle (GV)-to-metaphase II (MII) transition. In brief, *in vitro* transcribed eRF3a-ADAR2dd mRNAs or ADAR2dd-only mRNAs were microinjected into oocytes at the GV stage. Injected oocytes were maintained at the GV stage for 24 hours in the presence of milrinone (milrinone 24h) to allow sufficient translation of the injected mRNAs (Fig. 2c). For the MII stage, oocytes were released from the milrinone block after 12 hours and collected 12 hours later (milrinone 12h + release 12h) (Fig.2c). We collected four biological replicates for eRF3a-ADAR2dd mRNA-injected oocytes and two to three biological replicates for ADAR2dd-only mRNA-injected oocytes at each stage, which showed high correlation in RNA expression levels between replicates (Supplementary Fig. 5a, 5b). Furthermore, the RNA expression profiles of our samples correlated well with published datasets, supporting the high quality of our single-oocyte RNA-seq libraries (Supplementary Fig. 5a, 5b).

To assess RNA editing, we first compared EPKM values between replicates and found that eRF3a-ADAR2dd mRNA-injected oocytes showed substantially stronger inter-replicate correlation than ADAR2dd-only mRNA-injected oocytes (Fig. 2d, 2e and Supplementary Fig. 4a, 4b). Next, we compared the RNA editing profiles from our samples with published single-oocyte translatome data obtained using either the ligation-free, ultra-low-input, and enhanced Ribo-seq (Ribo-lite)^27^ or the combined RiboLace and SMARTer-seq approach (T&T-seq)^28^. As shown in Fig. 2f-i and Supplementary Fig. 5c-j, RNA editing profiles, represented by EPKM, in eRF3a-ADAR2dd mRNA-injected oocytes correlated higher with the translatome at both GV and MII stages than those in ADAR2dd-only controls. This observation is consistent with our findings in HEK293T cells, where RNA editing levels in eRF3a-ADAR2dd expressing cells correlated well with mRNA translation levels. Furthermore, the correlations between eRF3a-ADAR2dd EPKM and the translatome were comparable to those observed between different translatome datasets (Supplementary Fig. 5k-n). To further validate these results, we selected two representative genes (Hmgb2, and Mrpl10) that showed distinct translation dynamics across the two stages. Translation activity of Hmgb2 decreased at the MII stage (Supplementary Fig. 6a and 6g-i (the middle panel)), whereas Mrpl10 exhibited increased translation activity at the MII stage (Supplementary Fig. 6d and 6g-i (the left panel)), across all three datasets. Consistent with these observations, our immune-staining results revealed the corresponding changes in protein abundance, either reduced or increased fluorescence intensity, accompanied by opposite trends in RNA abundance from the GV to the MII stage (Supplementary Fig. 6b, 6c, 6e, and 6f). Additionally, we selected one gene (Rpl23) that exhibited inconsistent translation activity across three datasets for validation. Rpl23 showed higher translation activity, represented by EPKM, at the MII stage in our data (Fig. 2j). In contrast, it showed lower translation activity at the MII stage in both the Ribo-lite and T&T-seq datasets (Supplementary Fig. 6g-i). Our independent validation confirmed higher protein abundance and lower RNA abundance of Rpl23 at the MII stage (Fig. 2k, 2l). However, we also observed differences in the levels of differential RNA expression and translation activity between the two stages in our data compared to the Ribo-lite and T&T-seq data. These discrepancies may be attributed to differences in oocyte preparation strategies. Both Ribo-lite and T&T-seq used in vivo–collected immature and mature oocytes, whereas the oocytes analyzed in our method were matured in vitro. Together, our results demonstrate that, for Hmgb2, Rpl23, and Mrpl10, RAFTER-mediated RNA editing faithfully reflected translation dynamics across developmental stages. Therefore, RAFTER provides a reliable and feasible approach for measuring mRNA translation at the single-cell level.

In summary, we present a novel method, RAFTER, that integrates the RNA editing enzyme HyperTRIBE with the translation releasing factor eRF3a, enabling A-to-I editing during translation termination stage. We demonstrate that RAFTER not only exhibits specificity in detecting protein-coding mRNAs, but also shows sensitivity in identifying non-coding RNAs that encode small peptides. Furthermore, without requiring sophisticated experimental procedures, RAFTER achieves measurement accuracy comparable to both ribosome footprinting and polysome profiling. When combined with single-cell sequencing, RAFTER serves as an effective and accessible tool for profiling the translational landscape during mouse oocyte maturation. Together, these findings establish RAFTER as a simple yet powerful approach for monitoring translation with high accuracy, reproducibility, and broad applicability, even at the single-cell level.

### Online methods

#### Plasmid construction

To generate inducible expression plasmid for eRF3a-ADAR2dd fusion protein, the coding sequence of eRF3a (NM_002094.4_cds_NP_002085.3_1) was amplified from cDNA and ligated with the deaminase domain of human ADAR2 with a 25-amino-acid flexible linker in between. Additionally, an HA-T2A-mCherry cassette was inserted into the C-terminal end of the fusion protein. The resulting eRF3a-25aa linker-ADAR2dd-HA-T2A-mCherry cassette was cloned into the Tet-On 3G vector backbone to generate the doxycycline-inducible expression plasmid. Nucleotide sequences of fusion proteins are provided in Supplementary Table 1.

#### Cell culture and generation of cell lines

HEK293T cells were obtained from ATCC. Cells were cultured in Dulbecco’s Modified Eagle Medium (DMEM; Gibco, 11965092) supplemented with 10% fetal bovine serum (FBS; Cellmax, SA301.02) and 1% penicillin/streptomycin (P/S; Gibco, 15070063) at 37 °C in a humidified atmosphere containing 5% CO_2_.

RAFTER cell lines were generated using lentiviral transduction. Lentivirus were prepared using wiled type HEK293T cells at confluency about 70%. The target plasmid, pMD2.G plasmid, and psPAX2 plasmid were mixed at the ratio of 2:2:3 with polyethyleneimine (YEASEN, 40816ES03) in Opti-MEM (Gibco, 31985062). The mixture was incubated for 15 minutes at room temperature and then added to the cell culture medium. Two days later, the virus-containing medium was collected and mixed with polybrene (YEASEN, 40804ES76) for lentiviral transduction. Stably transduced cells were selected using 10 µg/mL blasticidin S (YEASEN, 60218ES10) for 5 days. Surviving cells were expanded for storage and further experiments.

#### Induction of RNA editing

RAFTER cells were induced with 1 µg/mL doxycycline in DMEM (10% FBS, 1% P/S) for 48 hours, followed by fluorescence-activated cell sorting to enrich the successfully induced cells according to mCherry signal. Total RNAs were extracted using TRlzol Reagent (Invitrogen, 15596026) according to the manufacturer’s protocol. RNA concentration was measured using Equalbit RNA BR Assay Kit (Vazyme, EQ212-01), and 500 ng total RNA was used as input material for the mRNA-seq library construction (YEASEN, 12309ES96) following the manufacturer’s protocol.

#### Data processing for RNA-seq

mRNA-seq libraries were sequenced on the Illumina NovaSeq 6000 System in PE150 mode. Sequencing quality was assessed with FastQC (v0.11.9, https://www.bioinformatics.babraham.ac.uk/projects/fastqc/). Adapter sequences were removed using Cutadapt (v1.18) with parameters: -j 40 -m 20 -q 20. The reads were then filtered for those derived from repetitive elements using sequences from the UCSC Genome Browser (https://genome.ucsc.edu/cgi-bin/hgTables) and afterwards aligned to the hg38 reference genome with STAR (v2.7.9a). Uniquely mapped reads were quantified using featureCounts (v2.0.1) against Gencode annotations (v37, Ensembl 103) with the following parameters: -T 24 -s 2 -p -B -C -g gene_name -Q 30.

#### RNA editing analysis for RNA-seq

After aligning reads to the genome, BAM files were further processed using SAILOR (v1.2.0)^15^ to identify A-to-I edit sites across the hg38 reference genome. Edit sites with confidence value larger than 0.65 were retained. The retained edit sites were further filtered based on the coverage (coverage ≥ 10) and mutation frequency (frequency < 0.95). Subsequently, for protein-coding genes, sites annotated within the 5’ UTR, CDS, or 3’ UTR were extracted for further analysis. For non-coding genes, sites annotated within the exon were extracted for further analysis. To characterize the distribution of RNA editing activity around translation start and stop sites, editing sites located within −2800 bp of the start and +2800 bp of the stop codon of the longest protein-coding transcript for each gene were extracted. The region was further divided into 40 equally sized bins, and the number of editing sites within each bin was counted to generate a positional density profile. To obtain a smooth representation of the editing distribution, cubic spline interpolation was applied to the binned counts and plotted the resulting continuous density curves.

Transcripts Per Kilobase Million (TPM) for each gene was calculated by normalizing read counts for sequencing depth and gene length. Genes with a TPM > 1 in the sample were included in Editing Score (ES) analyses. To compare editing at the gene level between the cells expressing eRF3a-ADAR2dd protein and ADAR2dd-only, the ES for each gene was calculated as the ratio between the number of edited reads and that of total reads mapped to the gene. To identify genes that were significantly edited, Fisher’s exact test was applied to examine the statistical significance of the editing score. The maximum p-value across the comparison results for all replicates was used as the final p-value for each gene. Then, the final p-value was adjusted for multiple comparisons using the Benjamini & Hochberg procedure (BH). Genes with an FDR < 0.05 and a LFC of ES > 1 were considered as significantly edited.

To more accurately reflect the association of fusion proteins to RNAs, we introduced EPKM (Edits Per Kilobase per Million total edits) as follows:

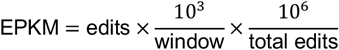

Specifically, the longest protein-coding transcripts of each gene was used for analysis. The window was limited from −100 to +2000 bp of the start codon and from −1000 to +200 bp of the stop codon. Edits represented the total number of editing events detected within the defined window of respective genes. Total Edits represented the sum of editing events across all genes of respective samples. Only genes with FPKM > 0 were considered.

#### In-vitro transcription

The EGFP-UTR cassette was cloned into the pcDNA6 plasmid, where a specific edit sites for eRF3a-ADAR2dd was in the UTR region. The resulting plasmids were linearized using NotI-HF and AgeI-HF (NEB, R3552). EGFP-UTR mRNAs were in-vitro transcribed using T7 High Yield RNA Synthesis Kit for Co-transcription (YEASEN, 10673ES50) from the linearized plasmid template. To generate mRNAs without the 5’ cap, the GAG component was substituted with RNase free H_2_O. In-vitro transcribed mRNAs were purified using RNA Clean & Concentrator-25 (ZYMO, R1018) and polyadenylated by *E*.*coli* poly(A) Polymerase (YEASEN, 14801ES60). To generate mRNAs without poly(A)-tail, the polyadenylation step was omitted. After polyadenylation, mRNAs were purified using RNA Clean & Concentrator-5 (ZYMO, R1014). Nucleotide sequences of fusion proteins are provided in Supplementary Table 2.

#### mRNA transfection and RNA editing quantification

RAFTER cells were seeded into 6-well plates at 0.4 million cells per well for 24 hours. Then 1 µg/mL doxycycline was supplemented into the culture medium for 36 hours. Next, 200 ng mRNAs were transfected into cells using mRNA Transfection Reagent (YEASEN, 40809ES01). Two hours after transfection, culture medium containing mRNAs was replenished by fresh culture medium containing 1 µg/mL doxycycline. Cells were collected for flow-cytometry analysis and RNA extraction after another 10 hours. Total RNAs were extracted using TRlzol Reagent according to the manufacturer’s protocol. RNA concentration was measured using Equalbit RNA BR Assay Kit and 2 µg total RNA was used for cDNA synthesis using Hifair III 1st Strand cDNA Synthesis Kit (gDNA digester plus). cDNA synthesis was conducted using random N6 primers only. Amplicons containing the edit site for Sanger sequencing were then amplified (Vazyme, P515) using primers shown in Supplementary Table 2. A-to-I editing level was quantified using the EditR package^29^. Primers for RT-qPCR were shown in Supplementary Table 2

#### Mouse oocytes preparation and injection

All animal procedures were approved by the Animal Research Committee of Nanjing Medical University and performed in accordance with Institutional guidelines. Fully grown GV stage oocytes were collected from 4-week-old female ICR mice. Before microinjection, oocytes were cultured in M2 medium (Sigma, M7167) with 2 µM milrinone (Sigma, M4659) to prevent germinal vesicle breakdown (GVBD). mRNAs encoding eRF3a-ADAR2dd or ADAR2dd-only were in vitro transcribed (HiScribe T7 ARCA mRNA Kit with tailing, NEB E2060S) and microinjected into oocytes (~5-10 pL at 500 ng/μL) using an Eppendorf Transforman 4R micromanipulator. After injection, oocytes were cultured for 24 hours under milrinone arrest to obtain GV stage samples. For MII stage samples, oocytes were cultured with milrinone for 12 hours, then released into milrinone-free medium for an additional 12 hours.

#### Immunofluorescence and confocal microscopy

For immunofluorescence staining, oocytes were fixed in 4% paraformaldehyde (in 1x PBS) for 30 min at room temperature, permeabilized in 0.5% Triton X-100 (in 1x PBS) for 20 min and blocked in 1% bovine serum albumin (in 1x PBS). Subsequently, oocytes were incubated overnight at 4 °C with primary antibodies diluted in blocking solution, followed by three washes in PBS and incubation with secondary antibodies for 1 hour at room temperature. Nuclei were counterstained with DAPI (5 μg/mL) for 15 min. After final washing, oocytes were mounted on glass slides using SlowFade Gold Antifade Reagent (Life Technologies) and imaged with a Zeiss LSM900 confocal microscope. Antibody details are provided in Supplementary Table 4. Semiquantitative analysis of fluorescence signals was performed using ImageJ software.

#### Single-oocyte RNA-seq

We constructed the single-oocyte RNA-seq libraries following the SMART-seq2 protocol^30^. Briefly, each oocyte was lysed in 2 μL of lysis buffer (0.2% Triton X-100 and 40 U/mL Recombinant RNase Inhibitor (Takara, 2313A)), then incubated in 10 μL Oligo(dT) buffer (1μL 10mM dNTP,1μL 10μM oligo(dT) Primer,8μL nuclease-free water) at 72 °C for 3 min and immediately placed on ice for 2 min. First-strand cDNA synthesis was performed by adding 4 μL of 5× RT buffer and 4 μL Reverse Transcriptase enzyme (ABclonal, RK20310), followed by cDNA amplification using Amplification Module PCR Mix (1 μL PCR primer and 29 μL PCR Master Mix) for 16 cycles. The resulting library was purified using 1× DNA Clean beads (Vazyme, N411-01). Tagmentation was carried out using Tn5 Enzyme mix and PCR mix (ABclonal, RK20237) to construct sequencing libraries from the amplified cDNA according to the manufacturer’s protocol. Final DNA library amplification was performed for 15 cycles using 25 μL PCR Mix and 5 μL amplification primers (10 μM), followed by purification with 0.6× and 0.15× DNA Clean beads (Vazyme, N411-01). Library quality was assessed using a PerkinElmer LabChip GX Touch system, and sequencing was performed on the Illumina NovaSeq 6000 platform in PE150 mode.

#### RNA isolation and real-time RT-qPCR

For each sample replicate, five oocytes were collected and lysed in 2 μL of lysis buffer (0.2% Triton X-100 and 40 U/mL Recombinant RNase Inhibitor (Takara, 2313A)), and cDNA synthesis was followed by retrotranscribing reverse transcription with primer transcript II reverse transcriptase (Takara, 2690A) according to the manufacturer’s protocol. RT-qPCR analysis was performed using Power SYBR Green PCR Master Mix (Vazyme, Q712-02) and an Applied Biosystems QuantStudio 5 Real-Time PCR system using the primers listed in Supplementary Table 3. Relative mRNA expression levels were calculated to the levels of endogenous 18S rRNA (used as a housekeeping gene), and each RT-qPCR experimental reaction was performed in triplicate.

#### Data processing for single-oocyte RNA-seq

Data processing for single-oocyte RNA-seq was performed similarly to the bulk RNA-seq analysis. Read alignment was conducted according to the GENCODE annotation (vM34, GRCm39), and SNP information was obtained from the dbSNP dataset (GVF_000001635.24, VCFv4.0). Strand orientation for each read was determined based on the GENCODE annotation. The resulting BAM files containing reads mapped to genes on the positive and negative strands were utilized as inputs to SAILOR (v1.2.0) for determining strand-specific A-to-I edit sites. For replicate comparisons, genes with EPKM or FPKM = 0 in all compared datasets were excluded. For the remaining genes, log2(EPKM+1) values were calculated, and Pearson correlation coefficients (R) were determined. Correlation plots were generated using the pandas.plotting module (v2.2.3).

#### Comparative analysis between RAFTER and analogous methods

Mean EPKM and FPKM were calculated across biological replicates within each group in our data. Ribo-seq data of HEK293T cells was obtained from Shu et al., 2025^21^. Polysome-seq data of HEK293T cells was obtained from Sun et al., 2024^22^. RPF and RNA data of the Ribo-lite approach were obtained from the Supplementary Table 1 of Xiong et al., 2022^27^. RPF and RNA data of the T&T-seq approach were obtained from the Supplementary Table of Hu et al., 2022^28^. To assess the relationship between RAFTER and analogous methods, genes with either EPKM or FPKM/TPM > 0 in all compared datasets were included. For RNA level comparison, respective values were transformed as log2(value+0.01) prior to calculating the Pearson correlation coefficient (R). For translation level comparison, respective values were transformed as log2(value+1) prior to calculating the Pearson correlation coefficient (R).

## Data Availability

The raw sequence data reported in this paper have been deposited in the National Genomics Data Center that are publicly accessible at https://ngdc.cncb.ac.cn/gsa-human/s/gd3658Zi, and https://ngdc.cncb.ac.cn/gsa/s/wwKF6Uq3. Source data are provided with this paper.

## Acknowledgments

This work was supported by The National Key R&D Program of China (Grant No. 2025YFC2708200 to X.Wang and 2022YFC3400400 to W.C.) and, Shenzhen Medical Research Fund (Grant No. B2502019 to W.C.), National Natural Science Foundation of China (Grant No. 32470590 to L.F., 92374117 to X.Wang and 32570659 to W.C.), The Shenzhen Science and Technology Program (Grant No. KQTD20180411143432337 to W.C.). We thank the members of the Chen lab for insightful discussions and suggestions during the process of this investigation. The authors acknowledge the Center for Computational Science and Engineering of SUSTech for the support on computational resources.

## Author contributions

W.C., X.Wang, L.F., and Z.S. conceived and designed the project. Z.S. performed the experiments with assistance from Y.L.. P.D. performed the computational analyses with assistance from X.Wen and F.J.. H.Y. collected the mouse oocytes and performed single-oocyte sequencing. W.C., X.Wang, L.F., and Z.S. wrote the manuscript with input from all authors.

## Figure legends

**Supplementary Figure 1. Expression of RAFTER in HEK293T cells. a-b**, Fluorescence activated cell sorting (FACS) results of HEK293T cells expressing ADAR2dd-only (**a**) or eRF3a-ADAR2dd (**b**). **c**, Western blot analysis of ADAR2dd-only and eRF3a-ADAR2dd protein levels in FACS sorted HEK293T cells. The experiment was repeated independently for 2 times with similar results.

**Supplementary Figure 2. RNA editing level in the 2**^**nd**^ **replicate. a-b**, Metaplots showing the distribution of confident edit sites (≥ 0.65 confidence score) in the 2^nd^ replicate. **a**, around stop codon. **b**, around start codon. **c-d**, Scatter plots showing the correlation of RNA editing levels between two replicates. **c**, in ADAR2dd-only cells. **d**, in eRF3a-ADAR2dd cells. The number of genes (n) and *Pearson* correlation coefficient (*R*) are indicated in the top left corner. **e-h**, Scatter plots showing the correlation between RNA editing levels and RNA translation levels. **e**, ADAR2dd-only (replicate 2) vs. Ribo-seq. **f**, eRF3a-ADAR2dd (replicate 2) vs. Ribo-seq. **g**, ADAR2dd-only (replicate 2) vs. Polysome-seq. **h**, eRF3a-ADAR2dd (replicate 2) vs. Polysome-seq. The number of genes (n) and *Pearson* correlation coefficient (*R*) are indicated in the top left corner. **i**, Scatter plots showing the correlation of RNA translation levels measured by Ribo-seq and Polysome-seq. The number of genes (n) and *Pearson* correlation coefficient (*R*) are indicated in the top left corner.

**Supplementary Figure 3. Expression of reporter mRNAs in HEK293T RAFTER cells. a-b**, Flow cytometry results of cells expressing reporter mRNAs. **a**, in ADAR2dd-only expressing cells. **b**, in eRF3a-ADAR2dd expressing cells. **c-d**, RT-qPCR quantification of RNA levels of reporter mRNAs. RNA levels were normalized by GAPDH. n = 3. **c**, in ADAR2dd-only expressing cells. **d**, in eRF3a-ADAR2dd expressing cells.

**Supplementary Figure 4. Correlation of RNA editing levels in ADAR2dd-only mouse oocytes. a-b**, Scatter plots showing the correlation of EPKM across ADAR2dd-only oocytes. The X and Y axes represent log2(EPKM+1). The number of genes (n) and *Pearson* correlation coefficient (*R*) are indicated in the top left corner. **a**, in ADAR2dd-only GV stage oocytes. **b**, in ADAR2dd-only MII stage oocytes.

**Supplementary Figure 5. Correlation of RNA levels and editing levels across mouse oocytes from different laboratories. a-b**, Scatter plots showing the correlation of RNA levels across oocytes from different laboratories. For RNA levels from this study and Ribo-lite approach, RNA levels are represented as log2(FPKM+0.01). For RNA levels from T&T-seq approach, RNA levels are represented as log2(TPM+0.01). The number of genes (n) and *Pearson* correlation coefficient (*R*) are indicated in the top left corner. **a**, GV stage oocytes. **b**, MII stage oocytes. **c-f**, Scatter plots showing the correlation between RNA editing levels measured by RAFTER in single mouse oocyte and RNA translation levels measured by the T&T-seq approach in 10 mouse oocytes. The X and Y axes represent RPF log2(TPM+0.01) and log2(EPKM+0.01), respectively. The number of genes (n) and *Pearson* correlation coefficient (*R*) are indicated in the top left corner. **c**, in GV stage oocytes, ADAR2dd-only vs. T&T-seq. **d**, in GV stage oocytes, eRF3a-ADAR2dd vs. T&T-seq. **e**, in MII stage oocytes, ADAR2dd-only vs. T&T-seq. **f**, in MII stage oocytes, eRF3a-ADAR2dd vs. T&T-seq. **g-j**, Scatter plots showing the correlation between RNA editing levels measured by RAFTER and RNA translation levels measured by the T&T-seq approach in single mouse oocyte. The X and Y axes represent RPF log2(TPM+1) and log2(EPKM+1), respectively. The number of genes (n) and *Pearson* correlation coefficient (*R*) are indicated in the top left corner. **g**, in GV stage oocytes, ADAR2dd-only vs. T&T-seq. **h**, in GV stage oocytes, eRF3a-ADAR2dd vs. T&T-seq. **i**, in MII stage oocytes, ADAR2dd-only vs. T&T-seq. **j**, in MII stage oocytes, eRF3a-ADAR2dd vs. T&T-seq. **k-n**, Scatter plots showing the correlation of RNA translation levels measured by the Ribo-lite and T&T-seq approach mouse oocytes. The X and Y axes represent T&T-seq RPF log2(TPM+1) and Ribo-lite RPF log2(FPKM+1), respectively. The number of genes (n) and *Pearson* correlation coefficient (*R*) are indicated in the top left corner. **k**, in GV stage oocytes, Ribo-lite singe oocyte vs. T&T-seq 10 oocytes. **l**, in GV stage oocytes, Ribo-lite singe oocyte vs. T&T-seq single oocyte. **m**, in MII stage oocytes, Ribo-lite singe oocyte vs. T&T-seq 10 oocytes. **n**, in MII stage oocytes, Ribo-lite singe oocyte vs. T&T-seq single oocyte.

**Supplementary Figure 6. Independent validation of differential RNA translation in mouse oocytes. a-c**, Hmgb2 results. **a**, RNA level and editing level of Hmgb2 detected in eRF3a-ADAR2dd mRNA-injected oocytes. n = 4. **b**, representative immunofluorescence results showing protein level of Hmgb2 in mouse oocyte. Scale bar, 20 μm. **c**, RT-qPCR quantification of RNA level (relative to endogenous 18S rRNA, n = 3) and fluorescence intensity quantification of protein level (from immunofluorescence results, n = 20). p (RT-qPCR) = 0.0002. p (fluorescence intensity) < 0.0001. Statistical significance was calculated using two-sided unpaired t test (default in Prism 9). The experiment was repeated independently for 2 times with similar results. **d-f**, Mrpl10 results. **d**, RNA level and editing level of Hmgb2 detected in eRF3a-ADAR2dd mRNA-injected oocytes. n = 4. **e**, representative immunofluorescence results showing protein level of Hmgb2 in mouse oocyte. Scale bar, 20 μm. **f**, RT-qPCR quantification of RNA level (relative to endogenous 18S rRNA, n = 3) and fluorescence intensity quantification of protein level (from immunofluorescence results, n = 19). p (RT-qPCR) < 0.0001. p (fluorescence intensity) = 0.0258. Statistical significance was calculated using two-sided unpaired t test (default in Prism 9). The experiment was repeated independently for 2 times with similar results. **g-i**, RNA level and RPF level of Rpl23, Hmgb2, and Mrpl10 measured by the Ribo-lite and T&T-seq approach, respectively. **g**, by the Ribo-lite approach in single oocyte. n =2. **h**, by the T&T-seq approach in single oocyte. n = 2. **i**, by the T&T-seq approach in 10 oocytes. n = 3.

## References

1. Buccitelli, C. & Selbach, M. mRNAs, proteins and the emerging principles of gene expression control. Nat Rev Genet 21, 630–644 (2020).

2. Truitt, M.L. & Ruggero, D. New frontiers in translational control of the cancer genome. Nat Rev Cancer 17, 332 (2017).

3. Kapur, M., Monaghan, C.E. & Ackerman, S.L. Regulation of mRNA Translation in Neurons-A Matter of Life and Death. Neuron 96, 616–637 (2017).

4. Tahmasebi, S., Khoutorsky, A., Mathews, M.B. & Sonenberg, N. Translation deregulation in human disease. Nat Rev Mol Cell Biol 19, 791–807 (2018).

5. Melamed, D. & Arava, Y. Genome-wide analysis of mRNA polysomal profiles with spotted DNA microarrays. Methods Enzymol 431, 177–201 (2007).

6. Ingolia, N.T., Ghaemmaghami, S., Newman, J.R. & Weissman, J.S. Genome-wide analysis in vivo of translation with nucleotide resolution using ribosome profiling. Science 324, 218–223 (2009).

7. Gingold, H. & Pilpel, Y. Determinants of translation efficiency and accuracy. Mol Syst Biol 7, 481 (2011).

8. McMahon, A.C. et al. TRIBE: Hijacking an RNA-Editing Enzyme to Identify Cell-Specific Targets of RNA-Binding Proteins. Cell 165, 742–753 (2016).

9. Brannan, K.W. et al. Robust single-cell discovery of RNA targets of RNA-binding proteins and ribosomes. Nat Methods 18, 507–519 (2021).

10. Xu, W., Abruzzi, K. & Rosbash, M. RiboTRIBE: Monitoring Translation with ADAR-meditated RNA Editing. bioRxiv (2021).

11. Jagannatha, P. et al. Long-read Ribo-STAMP simultaneously measures transcription and translation with isoform resolution. Genome Res 34, 2012–2024 (2024).

12. Frolova, L. et al. A highly conserved eukaryotic protein family possessing properties of polypeptide chain release factor. Nature 372, 701–703 (1994).

13. Frolova, L. et al. Eukaryotic polypeptide chain release factor eRF3 is an eRF1- and ribosome-dependent guanosine triphosphatase. RNA 2, 334–341 (1996).

14. des Georges, A. et al. Structure of the mammalian ribosomal pre-termination complex associated with eRF1.eRF3.GDPNP. Nucleic Acids Res 42, 3409–3418 (2014).

15. Deffit, S.N. et al. The C. elegans neural editome reveals an ADAR target mRNA required for proper chemotaxis. Elife 6 (2017).

16. Sun, Z. et al. Systematic characterization of the composition and dynamics of processing body-associated mRNAs. Nat Commun 16, 9867 (2025).

17. Martinez, T.F. et al. Accurate annotation of human protein-coding small open reading frames. Nat Chem Biol 16, 458–468 (2020).

18. van Heesch, S. et al. The Translational Landscape of the Human Heart. Cell 178, 242–260 e229 (2019).

19. Doll, S. et al. Region and cell-type resolved quantitative proteomic map of the human heart. Nat Commun 8, 1469 (2017).

20. Wells, S.E., Hillner, P.E., Vale, R.D. & Sachs, A.B. Circularization of mRNA by eukaryotic translation initiation factors. Mol Cell 2, 135–140 (1998).

21. Y. Shu et al. SPARP-seq reveals poly(A) tail shortening as a hallmark of the active human translatome. bioRxiv (2025).

22. Chen, W. et al. Systematic characterization of the composition and dynamics of processing body-associated mRNAs. Research Square (2024).

23. Chen, H. et al. Chemical and topological design of multicapped mRNA and capped circular RNA to augment translation. Nat Biotechnol 43, 1128–1143 (2025).

24. Chen, H. et al. Branched chemically modified poly(A) tails enhance the translation capacity of mRNA. Nat Biotechnol 43, 194–203 (2025).

25. Bennett, H.M., Stephenson, W., Rose, C.M. & Darmanis, S. Single-cell proteomics enabled by next-generation sequencing or mass spectrometry. Nat Methods 20, 363–374 (2023).

26. VanInsberghe, M., van den Berg, J., Andersson-Rolf, A., Clevers, H. & van Oudenaarden, A. Single-cell Ribo-seq reveals cell cycle-dependent translational pausing. Nature 597, 561–565 (2021).

27. Xiong, Z. et al. Ultrasensitive Ribo-seq reveals translational landscapes during mammalian oocyte-to-embryo transition and pre-implantation development. Nat Cell Biol 24, 968–980 (2022).

28. Hu, W. et al. Single-cell transcriptome and translatome dual-omics reveals potential mechanisms of human oocyte maturation. Nat Commun 13, 5114 (2022).

29. Kluesner, M.G. et al. EditR: A Method to Quantify Base Editing from Sanger Sequencing. CRISPR J 1, 239–250 (2018).

30. Picelli, S. et al. Smart-seq2 for sensitive full-length transcriptome profiling in single cells. Nat Methods 10, 1096–1098 (2013).

